# Activin Signaling Regulates Autophagy and Cardiac Aging through mTORC2

**DOI:** 10.1101/139360

**Authors:** Kai Chang, Ping Kang, Ying Liu, Kerui Huang, Erika Taylor, Antonia P. Sagona, Ioannis P. Nezis, Rolf Bodmer, Karen Ocorr, Hua Bai

## Abstract

Age-dependent loss of cardiac tissue homeostasis largely impacts heart performance and contributes significantly to cardiovascular diseases later in life. Cellular quality control machinery, such as autophagy/lysosome system, plays a crucial role in maintaining cardiac health and preventing age-induced cardiomyopathy and heart failure. However, how aging alters the autophagy/lysosome system to impact cardiac function remains largely unknown. Here using *Drosophila* heart as a model system, we show that activin signaling, a member of TGF-beta superfamily, negatively regulates cardiac autophagy and cardiac health during aging. We found that cardiac-specific knockdown of *Daw*, an activin-like protein in *Drosophila*, increased cardiac autophagy and prevented age-related cardiac dysfunction, including arrhythmia and bradycardia (slow heart rate). Inhibition of autophagy blocked *Daw* knockdown-mediated cardioprotection. Consistently, cardiac-specific expression of constitutively activated activin type I receptor *Babo* disrupted cardiac function at young ages. Intriguingly, the key autophagy regulator, mechanistic target of rapamycin complex 1 (mTORC1), was not involved in activin-mediated autophagy. Instead, activin signaling genetically interacted with Rictor, the key subunit of mTORC2, to regulate autophagy and cardiac aging. Knockdown of *Daw* increased the mRNA expression of Rictor and the phosphorylation of AKT in fly hearts. Finally, cardiac-specific silencing of *Daw* not only improved cardiac health, but also prolonged lifespan. Thus, our findings highlight an emerging role of activin signaling and mTORC2 in the regulation of autophagy and cardiac aging.

## Introduction

Aging is associated with an exponential increase of the incidence of cardiovascular diseases (CVD) ^1, 2^. Age-related changes in cardiovascular structure and output have been linked to increased risk of coronary heart disease, sudden cardiac death and stroke in elderly population ^3^. During normal aging, the left ventricular wall of human hearts becomes thickened and the diastolic filling rate of left ventricle gradually decreases. On the other hand, the left ventricular systolic performance at rest remains no change with age ^3^. Several mechanisms underlying these age-associated changes in cardiovascular structure and function have been proposed, for example changes in growth factor signaling, decreased cellular quality control, altered calcium handling, elevated extracellular matrix deposition or fibrosis, increased mitochondria damage, and the production of reactive oxygen species (ROS) ^2, 4^. Resolving the contributing mechanisms underlying age-dependent decline of cardiovascular function is critical for the development of therapeutic interventions for the treatment of cardiovascular diseases.

Cellular quality control systems, such as macroautophagy (hereinafter referred to as autophagy), are essential to the maintenance of tissue homeostasis during aging ^5, 6^. Autophagy is a highly conserved process that is responsible for the degradation and recycling of damaged organelles, protein aggregates and other cytoplasmic substances. The autophagic process initiated with the isolated membrane, or phagophore that elongates to form a double-membrane structure, the autophagosome. Then the autophagosome fuses with a lysosome to form an autolysosome to degrade the cargo within the autolysosome ^7^. It is generally accepted that autophagy activity declines with age ^8-10^. Disruption of autophagy pathways often leads to loss of tissue homeostasis and tissue dysfunction. For example, knocking out autophagy gene Atg5 in the mouse heart accelerates cardiac aging, including an increase in left ventricular hypertrophy and decrease in factional shortening ^10^. As a crucial autophagy regulator, mechanistic target of rapamycin (mTOR), especially mTOR complex 1 (mTORC1), plays an important role in the regulation of cardiac tissue homeostasis ^2^. Rapamycin-mediated suppression of mTORC1 induces autophagy and protects mouse cardiomyocytes from oxidative stress ^11^. In *Drosophila*, mTORC1 is known to promote electrical pacing-induced heart failure ^12, 13^, and high-fat-diet-induced cardiomyopathy ^14^. The mTOR signaling consists of two complexes, mTORC1 and mTORC2. It is not known whether mTORC2 also contributes to the regulation of autophagy ^15^. Although inhibition of mTOR is one of the robust longevity interventions ^16, 17^, it remains unclear whether mTOR pathway plays a major role in age-dependent decline of autophagy. Notably, a recent study showed that mTORC1 activity does not increase with age in most tissues of C57BL/6J.Nia mice ^18^. Thus, the mechanistic basis for age-dependent alterations in autophagy and the contribution of mTORC1 remains to be determined.

Autophagy can be activated by various stresses, such as dietary restriction, hypoxia, oxidative stress, and pathogen infection ^19^. The activation of autophagy is tightly controlled by key cellular signaling pathways, in particular the nutrient signaling pathways like mTORC1 and AMP-activated protein kinase (AMPK) ^20^. These pathways are often involved in control of longevity. We and other groups recently found that activin signaling, a transforming growth factor beta (TGF-beta) subfamily, regulates muscle proteostasis and longevity in *Drosophila* ^21, 22^. As in mammals, TGF-beta family in *Drosophila* has two major branches, bone morphogenetic protein (BMP) and activin signaling pathways ^23^. In both pathways, signaling starts with ligand binding to a receptor complex composed of type I and type II receptor kinases, followed by phosphorylation of receptor-activated Smad (R-Smad). *Drosophila* activin signaling is known to regulate neuroblast proliferation, axon guidance, and metabolic homeostasis during development (reviewed in ^24^). The regulation of adult physiology by *Drosophila* activin is not well understood. We previously showed that knockdown of one of the three activin ligands, Daw, prolongs lifespan and increases autophagosome formation in flies ^21^. However, the role of TGF-beta/activin signaling in cardiac aging remains largely unknown.

It has been shown that TGF-beta factors promote cardiac fibrosis during aging ^25^. Mice with heterozygous mutants for activin receptor-like kinases (ALK4) are protected from pressure overload-induced fibrosis and dysfunction ^26, 27^. Inhibition of ALK5 also reduces myocardial infarction-induced systolic dysfunction and left ventricular remodeling in rats ^28^. However, conflicting results are reported in other studies showing that activin A protects hearts from hypoxia/reoxygenation- and ischemia/reperfusion-induced cell death ^29^. In addition, it is unknown whether TGF-beta family proteins, in particular activin, regulate cardiac homeostasis and function through autophagy. Given that the cardiac development and aging are remarkably conserved between *Drosophila* and mammals, we investigated the role of activin signaling in the regulation of cardiac autophagy and cardiac aging using *Drosophila* heart model and cardiac movement detection techniques ^30^. Here we show that activin signaling acts on hearts to regulate autophagy and age-related cardiomyopathy in *Drosophila*. RNAi-mediated knockdown of the activin-like protein *Daw* preserved cardiac function with age, while expression of activated activin type I receptor *Babo* induced cardiomyopathy early in life. Inhibition of autophagy blocked the cardioprotective effects by *Daw* knockdown. Intriguingly, activin signaling genetically interacted with mTORC2 subunit Rictor (rapamycin-insensitive companion of mTOR), but not mTORC1 to control autophagy and cardiac aging. Thus, our study uncovered a novel interaction between activin and mTORC2 in the regulation of tissue homeostasis, autophagy, and age-related cardiomyopathy.

## Results

### Heart-specific knockdown of activin-like ligand *Daw* slows cardiac aging

Our previous study demonstrated that the activin-like protein Daw regulated tissue homeostasis (especially in flight muscle) and longevity in *Drosophila* ^21^. Interestingly, we found that activin signaling increased in aging fly hearts, as indicated by elevated phosphorylation of Smad2 in heart nuclei (Fig. 1a). This finding suggest that activin signaling may play a role in regulating cardiac aging. To investigate this possibility, we knocked down the expression of *Daw* specifically in the heart with a binary GAL4/UAS system where we crossed *UAS-Daw RNAi* lines into a cardiac-specific tissue driver (*Hand-gal4*). We analyzed cardiac performance in young and old flies using a high-speed video imaging system (semi-automatic optical heartbeat analysis, SOHA) ^31^. M-Mode traces from the SOHA analysis indicated that knockdown of *Daw* preserved cardiac contractility at advanced ages (Fig. 1b). In wild-type flies, normal aging is associated with increased cardiac arrhythmia, diastolic intervals, heart period (bradycardia, slow heart rate) (Figs. 1c, 1d, 1e), and decreased cardiac output (Fig. 1f). Systolic intervals normally remain unchanged (Fig. 1g). Interestingly, cardiac-specific knockdown of *Daw* attenuated age-dependent increase in arrhythmia, diastolic intervals, and heart period (Figs. 1c, 1d, 1e). *Daw* knockdown also maintained relatively normal cardiac output at advanced ages (Fig. 1f). Similar results were observed from three independent Daw RNAi lines (the knockdown efficiency of Daw RNAi was verified by qRT-PCR in our previous studies ^21^).

**Figure 1.**
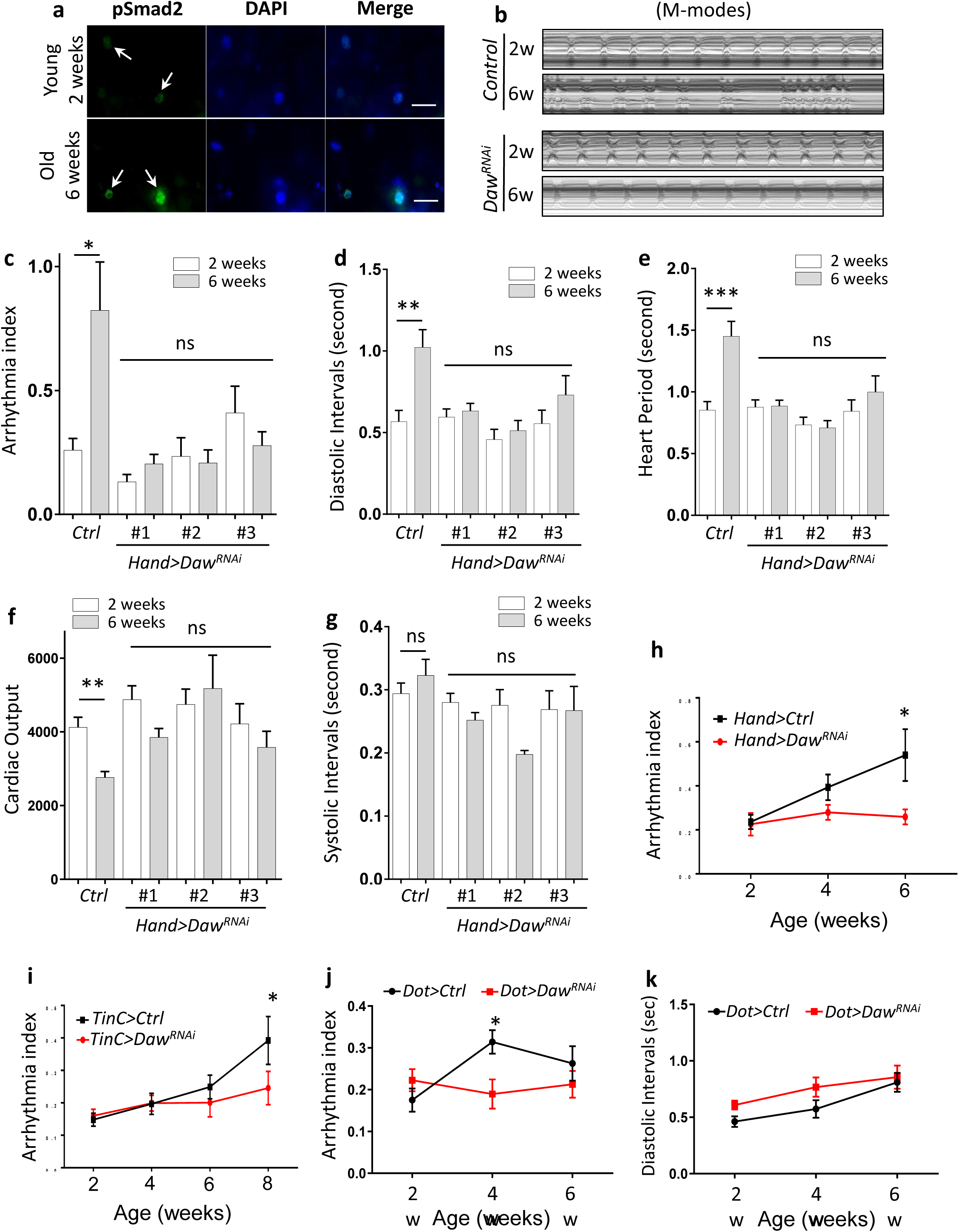
Heart-specific knockdown of *Daw* slows cardiac aging. **a.** Immunostaining of phospho-Smad2 in fly hearts at young (2 weeks) and old ages (6 weeks). Arrows indicate cardiomyocyte nuclei and positive pSmad2 staining. Scale bar: 20 μm. **b.** Representative M-mode traces (8 seconds) showing age-dependent movement of heart wall in control and heart-specific *Daw* knockdown flies (*Hand-gal4>UAS-Daw*^*RNAi*^). **c-g.** Age-related changes in arrhythmia index, diastolic intervals, heart period, cardiac output, and systolic intervals in control (*Ctrl*) and cardiac-specific *Daw* knockdown flies (*Daw* ^*RNAi*^). Flies were cultured at 40% relative humidity. *Hand-gal4* driver was used to knock down gene expression specifically in cardiac tissues (cardiomyocytes and pericardial cells). Results from three independent *UAS-Daw*^*RNAi*^ lines are shown (RNAi #1: BDSC #34974, RNAi #2: VDRC #105309, RNAi #3: BDSC #50911). Statistical significance is assessed by two-way ANOVA followed by Tukey multiple comparisons test (* p<0.05, ** p<0.01, *** p<0.001, ns = not significant). N=7~20. The interaction between genotype and age is statistically significant for heart period (P = 0.0041) and diastolic interval (P = 0.0243). **h-j.** Age-dependent changes in cardiac arrhythmia between control and *Daw* knockdown using various tissue drivers, *Hand-gal4* (cardiomyocytes and pericardial cells), *TinC-gal4* (cardiomyocytes), and *Dot-gal4* (pericardial cells). Student t-test (* p<0.05, ** p<0.01, *** p<0.001). N=25~31. **k.** Age-dependent changes in diastolic intervals between control and *Daw* knockdown in pericardial cells *Dot-gal4*). Student t-test (* p<0.05, ** p<0.01, *** p<0.001). N=13~26.

The *Hand-gal4* line drives gene expression in both cardiomyocytes and pericardial cells ^32^ To test the tissue-specific role of Daw in cardiac aging, we first crossed Daw RNAi into a cardiomyocyte-specific driver *TinC-gal4* ^33^. Similar to the results from *Hand-gal4* (Fig. 1h), cardiomyocyte-specific knockdown of *Daw* prevented age-related increase in cardiac arrhythmia (Fig. 1i). We also tested whether *Daw* expression in pericardial cells can regulate cardiac aging using a pericardial cell-specific driver *Dot-gal4* ^34^. Interestingly, knockdown of *Daw* in pericardial cells also attenuated age-induced arrhythmia (Fig. 1j), but not the increase in diastolic interval (Fig. 1k). Thus, Daw appears to promote cardiac aging, in particular age-induced arrhythmia, through both cardiomyocyte-specific regulation and non-autonomous signaling from pericardial cells.

### Cardiomyocyte-specific knockdown of activin receptor *Babo* delays cardiac aging

In *Drosophila*, activin-like ligand Daw signals through the type I receptor Babo, the fly homolog of mammalian ALK4 (activin receptor type-1B) ^23^. To investigate whether Daw acts through its receptor Babo to modulate cardiac aging, we first asked if Babo plays a similar role in preventing age-dependent increase in cardiomyopathy. Consistent with the results from *Daw* knockdown, we found that cardiomyocyte-specific knockdown of *Babo* blocked the age-related increase in cardiac arrhythmia, diastolic intervals, and heart period (Figs. 2a, 2b, 2c). Two independent Babo RNAi lines were used and both gave similar results. Systolic intervals remained unchanged in both controls and *Babo* RNAi flies (Fig. 2d). Knockdown of *Babo* did not affect age-dependent changes in cardiac output (Data not shown).

**Figure 2.**
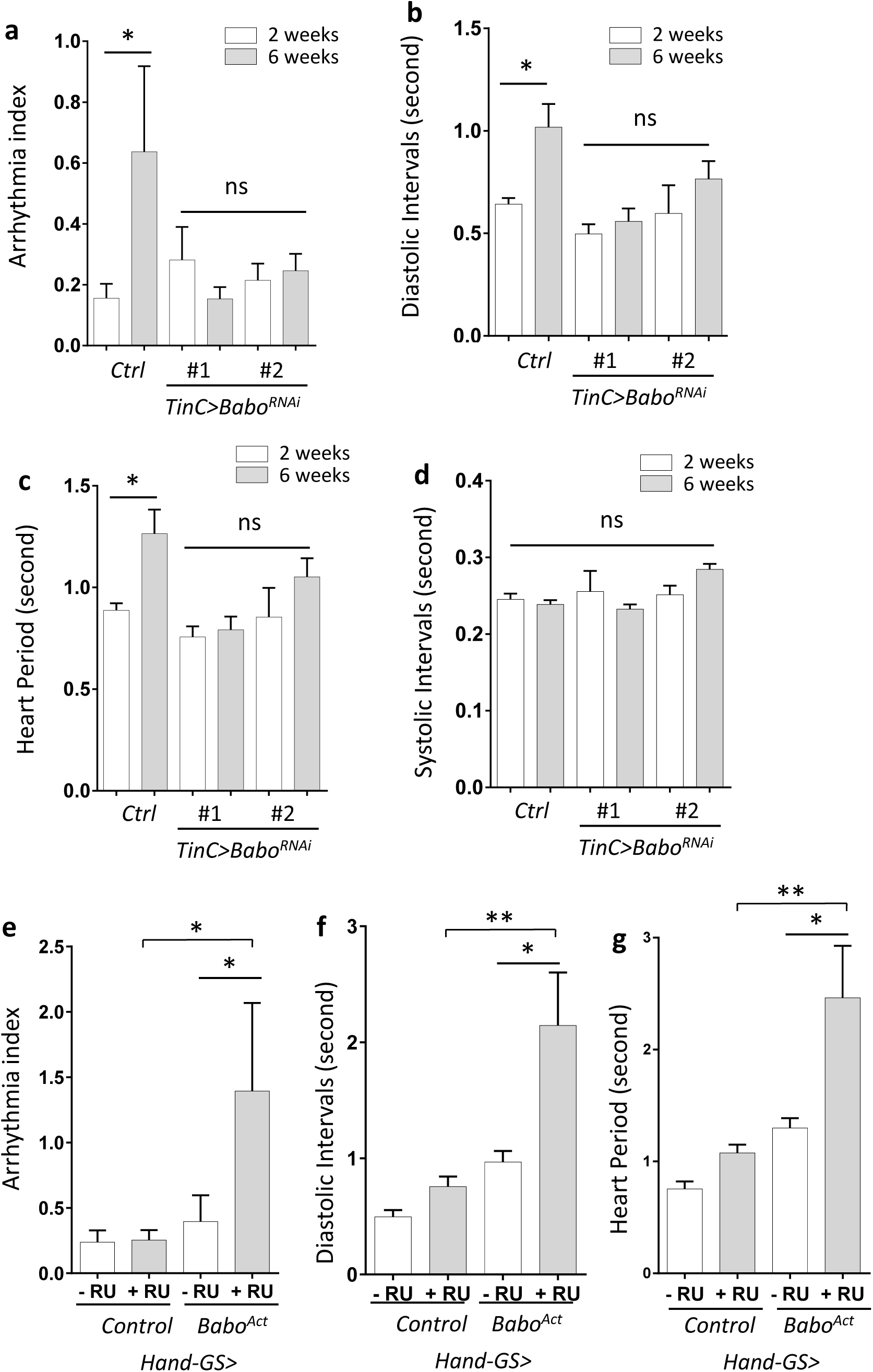
Cardiomyocyte-specific knockdown of activin receptor *Babo* delays cardiac aging. **a-d.** Age-dependent changes in cardiac arrhythmia, diastolic intervals, heart period, and systolic intervals in control (*Ctrl*) and cardiomyocyte-specific *Babo* knockdown flies (*Babo*^*RNAi*^). Flies were cultured at 40% relative humidity. *TinC-gal4* driver was used. Results from two independent *UAS-Babo*^*RNAi*^ lines are shown (RNAi #1: BDSC #25933, RNAi #2: BDSC #40866). N=15~30. **e-g.** Changes in cardiac arrhythmia, diastolic intervals and heart period in fly hearts expressing constitutively activated *Babo* (*Babo*^*Act*^). GeneSwitch heart driver *Hand-GS-gal4* was used to induce adult-onset *Babo* activation. RU: RU486 (Mifepristone). Flies were cultured at 40% relative humidity. N=6~9. One-way ANOVA (*** p<0.001, ** p<0.01, * p<0.05, ns = not significant).

In a reciprocal experiment, we used a GeneSwitch heart driver (*Hand-GS-gal4*) to express a constitutive active form of *Babo* (*Babo-Act*, *or Babo.Q302D*) specifically in adult hearts. We found that cardiac expression of activated *Babo* significantly increased cardiac arrhythmia, diastolic intervals, and heart period already at young ages (Figs. 2e, 2f, 2g). Taken together, these results suggest that Babo plays an important role in age-related cardiomyopathy, especially cardiac arrhythmia, diastolic function and heart rate.

### Daw negatively regulates autophagy in fly hearts

Autophagy is one of the key mechanisms in maintaining tissue hemostasis, and its activity normally declines with age ^6, 9, 10^. Our previous studies showed that activin signaling regulates autophagosome formation and Atg8a transcription ^21^. It is likely that Daw regulates cardiac aging by modulating autophagy. To first verify the role of Daw in autophagy-lysosome activity, we used LysoTracker staining to monitor the lysosomal acidification in adult fat body, a well-established autophagy marker in flies ^35^. Consistent with our previous finding, heterozygous

*Daw*^*[11]*^ loss-of-function mutants showed increased LysoTracker staining, suggesting an elevated basal autophagy/lysosome activities (Fig. 3a). Using a lipidation assay, we furthermore showed that mutation of *Daw* increased the levels of lipidated Atg8a, the fly homolog of mammalian LC3/GABARAP family proteins (Fig. 3b). Finally, mosaic analysis revealed that somatic cell clones in larval fat body expressing RNAi against either Daw or Babo (marked by GFP) showed increased numbers of autophagosome, indicated by mCherry-Atg8a puncta (Fig. 3c). The data suggest that activin signaling negatively regulates autophagy and lysosome activities.

**Figure 3.**
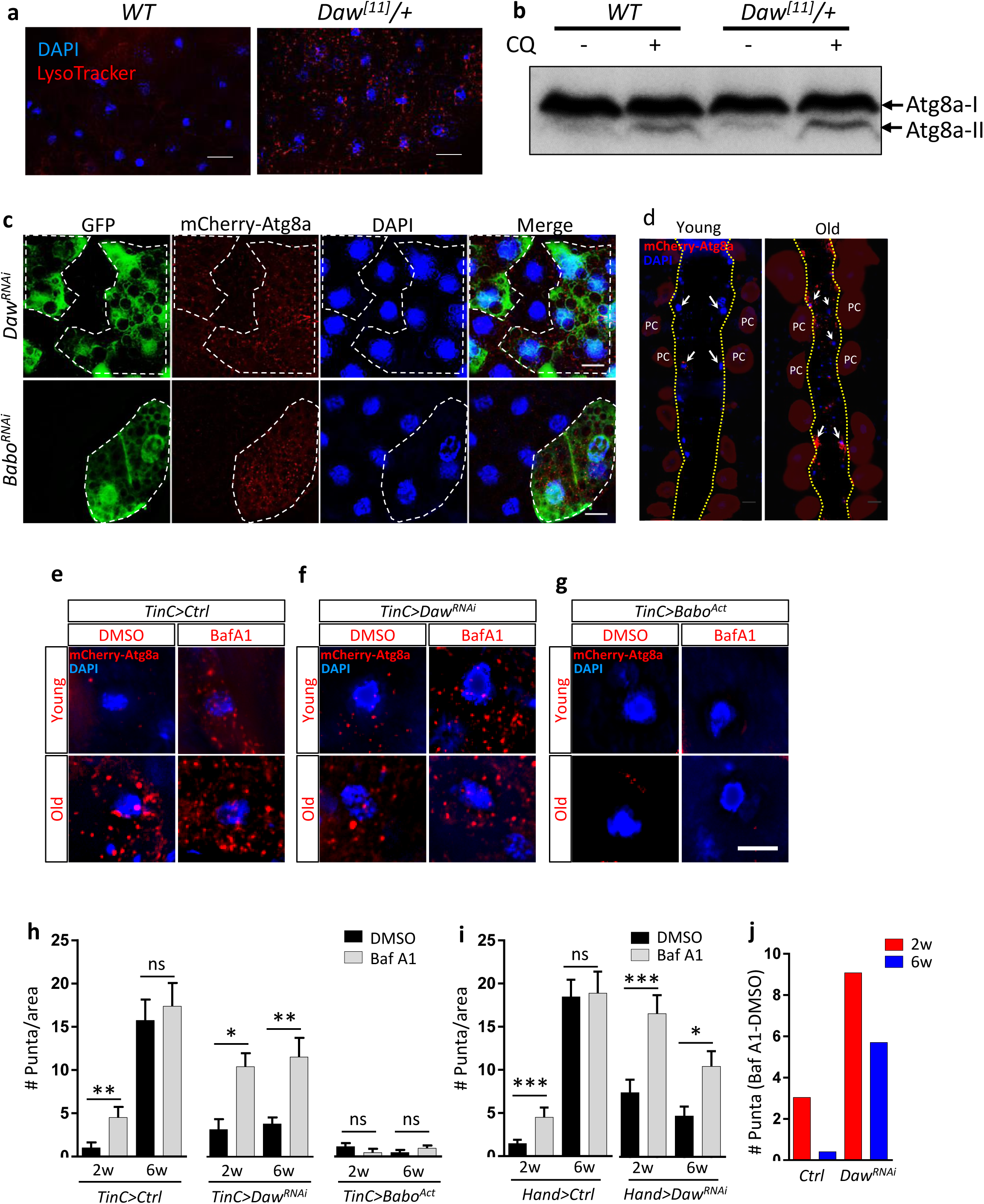
Daw negatively regulates autophagy in fly hearts. **a.** Representative images of LysoTracker staining of young adult fat body between wild-type (*WT*) and heterozygous *Daw*^*[11]*^ mutants. Scale bar: 20 μm. **b.** Western blot analysis on Atg8a lipidation between *WT* and *Daw*^*[11]*^ mutants. GABARAP antibodies were used to detect Atg8a. The lower band represents lipidated Atg8a. Flies were fed with chloroquine (CQ, 20 mM) for at least 24 hours prior to western blots. **c.** Representative images of mosaic analysis on autophagosome staining in larval fat body. Fat body clones were generated by crossing *Daw*^*RNAi*^ and *Babo*^*RNAi*^ lines into FLPout line carrying mCherry-Atg8a reporter (*hs-flp; endogenous P-3x mCherry-Atg8a*, *UAS-GFP/Cyo;Act>CD2>Gal4,UAS-Dcr2*). RNAi Clones are GFP-positive cells (dashed lines). Scale bar: 20 μm. **d.** Representative images of fly hearts expressing autophagosome reporter mCherry-Atg8a at both young and old ages. Arrows indicate cardiomyocyte nuclei and surrounding autophagosomes. Abdominal segments A2-A3 are shown. Heart tube is located between two yellow dashed lines. PC: Pericardial cell. **e-g.** Representative images of mCherry-Atg8a reporter surrounding cardiomyocyte nuclei in control, *Daw*^*RNAi*^, and *Babo*^*Act*^ flies. Semi-intact hearts were incubated with 100 nM of bafilomycin A1 (Baf A1) for two hours prior to immunostaining. Scale bar: 10 μm. **h.** Quantification of age-dependent changes in mCherry-Atg8a-positive puncta in perinuclear region in control, *Daw*^*RNAi*^, and *Babo*^*Act*^ flies. *TinC-gal4* was used to drive the expression of Atg8a reporter and gene knockdown. One-way ANOVA (*** p<0.001, ** p<0.01, * p<0.05, ns = not significant). N=5~7. **i.** Quantification of age-dependent changes in mCherry-Atg8a-positive puncta in perinuclear region in control and *Daw*^*RNAi*^. *Hand-gal4* was used to drive the expression of Atg8a reporter and *Daw* knockdown. One-way ANOVA (*** p<0.001, ** p<0.01, * p<0.05, ns = not significant). N=5~7. **j.** Quantification of Baf A1-induced mCherry-Atg8a-positive puncta between control and *Daw*^*RNAi*^.

Next, we examined whether Daw plays a similar role in modulating autophagy in fly hearts. Using a commercial antibody for *Drosophila* Atg8a protein (Fig. S1 shows the antibody validation), we found that the number of autophagosome in the heart was higher in *Daw*^*[11]*^ mutants compared to wild-type (Fig. S2a). Because autophagy is a dynamic process, the increased autophagosome number could result from increased autophagy activities, or blockage of lysosomal degradation ^36^. We next examined the role of Daw in the regulation of autophagic flux (the turnover of autophagosome). We exposed semi-intact fly hearts to a V-ATPase inhibitor bafilomycin A1 (Baf A1), which blocks both lysosomal acidification and autophagosome-lysosome fusion ^37^. We found that Baf A1 treatment resulted in a higher increase in the number of Atg8a-positive autophagosome in *Daw*^*[11]*^ hearts compared to wild-type hearts (Figs. S2a-S2c). Thus, the results suggest that Daw negatively regulate autophagic flux in fly hearts.

Aging tissues show reduced autophagy and accumulated cellular damages. We next examined whether inhibition of activin/Daw signaling in fly hearts can rescue age-dependent decline of autophagy. Using a mCherry-Atg8a reporter line, we observed a dramatic accumulation of Atg8a-positive autophagosome in the heart (Fig. 3d, 3e, 3h). The cardiomyocyte-specific driver *TinC-gal4* was used. The accumulation of autophagosome is likely due to the blockage of autophagosome turnover, because increases numbers of Atg8a-positive puncta were only seen in Baf A1-treated young hearts, but not the old hearts (Figs. 3e, 3h, 3j). Consistently, the adaptor protein p62/Ref(2)P, a marker for defective aggrephagy ^38^, was accumulated in aging hearts and Atg8a mutants (Fig. S3). Intriguingly, flies with heart-specific *Daw* knockdown maintained high levels of cardiac autophagosome turnover (or active autophagic flux) at both young and old ages, as indicated by the induction of autophagosome number upon Baf A1 treatment (Figs. 3f, 3h, 3j). Notably, constitutively activated Babo repressed the autophagic flux at both young and old ages (Figs. 3g, 3h). Activation of Babo also blocked the accumulation of autophagosome in old fly hearts, which suggests that besides the regulation of autophagosome turnover, activin signaling also play an important role in autophagosome initiation and formation (Figs. 3g, 3h). We were able to verify these phenotypes using another heart-specific driver *Hand-Gal4* (Fig. 3i). Taken together, our findings suggest that inhibition of activin/Daw signaling promotes autophagosome turnover and maintains healthy autophagic activity in old hearts.

### Inhibition of autophagy, but not activation of mTORC1, blocks *Daw* knockdown-mediated cardioprotection

Since we observed an induction of autophagy in *Daw* knockdown flies, we next asked whether autophagy activity is required for Daw-regulated cardiac aging. First, we fed control and *Daw* knockdown flies with lysosomal inhibitor chloroquine (CQ). As expected, CQ treatment increased cardiac arrhythmia in *Daw* knockdown flies at old age (Fig. 4a). To directly test whether autophagy plays any role in Daw-regulated cardiac aging, we generated double knockdown flies by combining *UAS-Daw RNAi* and *UAS-Atg1 RNAi* fly lines. Atg1 is a serine/threonine-protein kinase that is essential for autophagy initiation. Cardiac-specific knockdown of *Atg1* increased both cardiac arrhythmia and diastolic intervals at young ages (Fig. 4b), similar to the fly heart with activated Babo (Figs. 2e, 2f). Consistent with CQ treatment, cardiac-specific knockdown of *Atg1* attenuated cardioprotective effects of Daw RNAi, since simultaneously knockdown of *Atg1* and *Daw* in the heart led to age-dependent increase in arrhythmia similar to control flies (Fig. 4c). Thus, these results suggest that autophagy and Atg1 are required for Daw-regulated cardiac aging.

**Figure 4.**
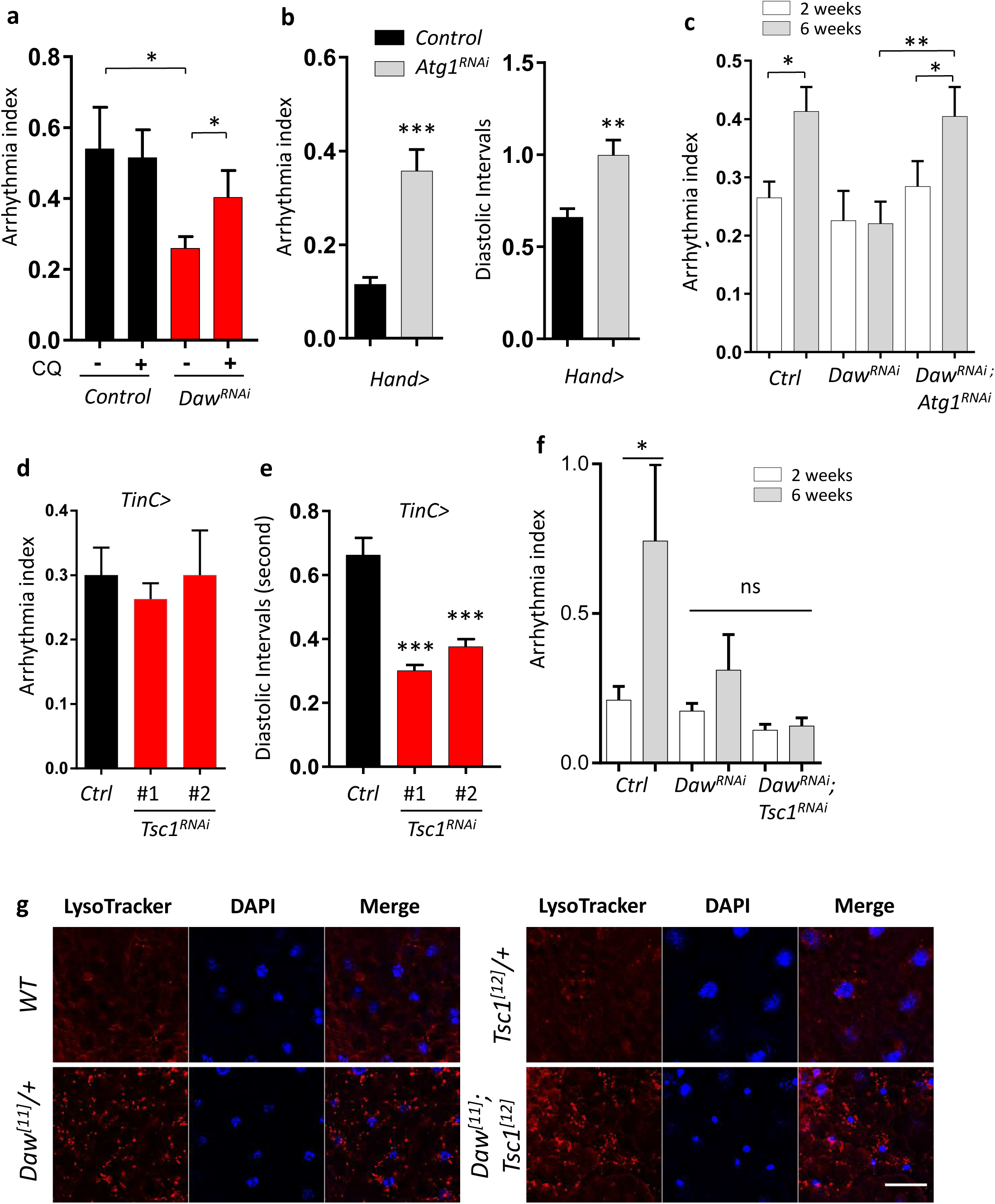
Inhibition of autophagy, but not activation of mTORC1, blocks *Daw* knockdownmediated cardioprotection. **a.** Cardiac arrhythmia of chloroquine-treated (20 mM, 24 hours) 6-week-old control and *Daw* knockdown flies (*Hand-gal4* used). One-way ANOVA (*** p<0.001, ** p<0.01, * p<0.05, ns = not significant). N=14~38. **b.** Cardiac arrhythmia and diastolic intervals in heart-specific knockdown of *Atg1* at young ages (2-week-old) (*Hand-gal4* used). One-way ANOVA (*** p<0.001, ** p<0.01, * p<0.05, ns = not significant). N=16~19. **c.** Age-dependent changes in cardiac arrhythmia in control, *Daw*^*RNAi*^, and *Daw*^*RNAi*^; *Atg1*^*RNAi*^ flies (*TinC-gal4* used). One-way ANOVA (*** p<0.001, ** p<0.01, * p<0.05, ns = not significant). N=16~35. **d-e.** Cardiac arrhythmia and diastolic intervals of *Tsc1* knockdown flies (*TinC-gal4* used). Two independent Tsc1 RNAi lines were used (RNAi-1: BDSC #52931, RNAi-2: BDSC #54034). One-way ANOVA (*** p<0.001, ** p<0.01, * p<0.05, ns = not significant). N=16~18. **f.** Age-dependent changes in cardiac arrhythmia in control, *Daw*^*RNAi*^, and *Daw*^*RNAi*^; *Tsc1*^*RNAi*^ flies (*TinC-gal4* used). One-way ANOVA (*** p<0.001, ** p<0.01, * p<0.05, ns = not significant). N=7~30. **g.** Representative images of LysoTracker staining in adult fat body of *WT*, *Daw*^*[11]*^, *Tsc1*^*[12]*^ and double mutant *Daw*^*[11]*^; *Tsc1*^*[12]*^. Scale bar: 10 μm.

mTORC1 is a key negative regulator of autophagy ^35^. We then tested whether Daw could negatively regulate autophagy and cardiac aging through activation of mTORC1 signaling. To test this possibility, we first generated double knockdown flies by combining *UAS-Daw RNAi* and *UAS-Tsc1 RNAi*. Tsc1 (Tuberous sclerosis protein 1) is a negative regulator of mTORC1 (Fig. S4a). Surprisingly, activation of mTORC1 via *Tsc1* knockdown had no effects on cardiac arrhythmia (Fig. 4d), but reduced diastolic intervals at young age (Fig. 4e), which is very different from the cardiomyopathy resulted from cardiac-specific expression of activated activin receptor *Babo* (Figs. 2e, 2f). Furthermore, knockdown of *Tsc1* did not rescue the cardioprotective effects of *Daw* knockdown during aging (Fig. 4f). Simultaneously knockdown of *Tsc1* and *Daw* did not show age-dependent increase in arrhythmia. Similarly, loss-of-function mutant *Tsc1*^*[12]*^ did not rescue the elevated LysoTracker staining in *Daw*^*[11]*^ (Fig. 4g). These results suggest that activation of mTORC1 is not required for Daw-regulated autophagy and cardiac aging.

It is known that inhibition of mTORC1 prolongs lifespan ^16, 39^ and attenuates stress-induced cardiomyopathies ^12, 13, 40, 41^. However, the effects of mTORC1 induction on cardiac aging are not well understood. Here, we observed unique changes of cardiac function in response to activation of mTORC1 (via *Tsc1* knockdown), including decreased heart period (Tachycardia, fast heart rate), and reduced diastolic intervals (Figs. 4d, 4e) These cardiac functional changes are not the same as age-induced cardiomyopathies, rather they resemble high-fat-diet-induced cardiac complications where mTORC1 is known to play an important role ^14^. Intriguingly, flies with cardiac-specific knockdown of *Tsc1* showed no age-dependent increases in heart period, diastolic intervals, and arrhythmia (Figs. S4b, S4c, S4d). These unique alterations in cardiac function were also observed in other mTORC1 activation mutants, such as 1) Knockdown of *Reptor* (Repressed by TOR), a recently identified bZIP transcription factor that is negatively regulated by mTORC1 ^42^ (Figs. S4e, S4f, S4g), and 2) Over-expression of *Rheb* (Ras Homolog Enriched in Brain), a positive regulator of mTORC1 (Figs. S4h, S4i, S4j).

### Daw genetically interacts with mTORC2/Rictor to control autophagy and cardiac aging

Although Daw did not interact mTORC1 to regulate autophagy and cardiac aging, cardiac-specific knockdown of *Daw* induced the phosphorylation of AKT, the upstream regulator of mTORC1 (Figs. 5a, 5b). On the other hand, activated receptor Babo reduced cardiac AKT phosphorylation (Figs. 5c, 5d). It is known that AKT can be activated and phosphorylated by two kinases at distinct positions, PDK1 (phosphoinositide-dependent kinase 1) at Thr308 and Rictor (major component of mTORC2) at Ser473 (Ser505 in flies) ^43^. Cardiac-specific over-expression Rictor can also induce AKT phosphorylation (Figs. 5e, 5f). Interestingly, knockdown of *Daw* up-regulated the mRNA expression of *Rictor* in fly hearts (Figs. 5g, 5h). These results suggest that Daw is a negative regulator of mTORC2/Rictor.

**Figure 5.**
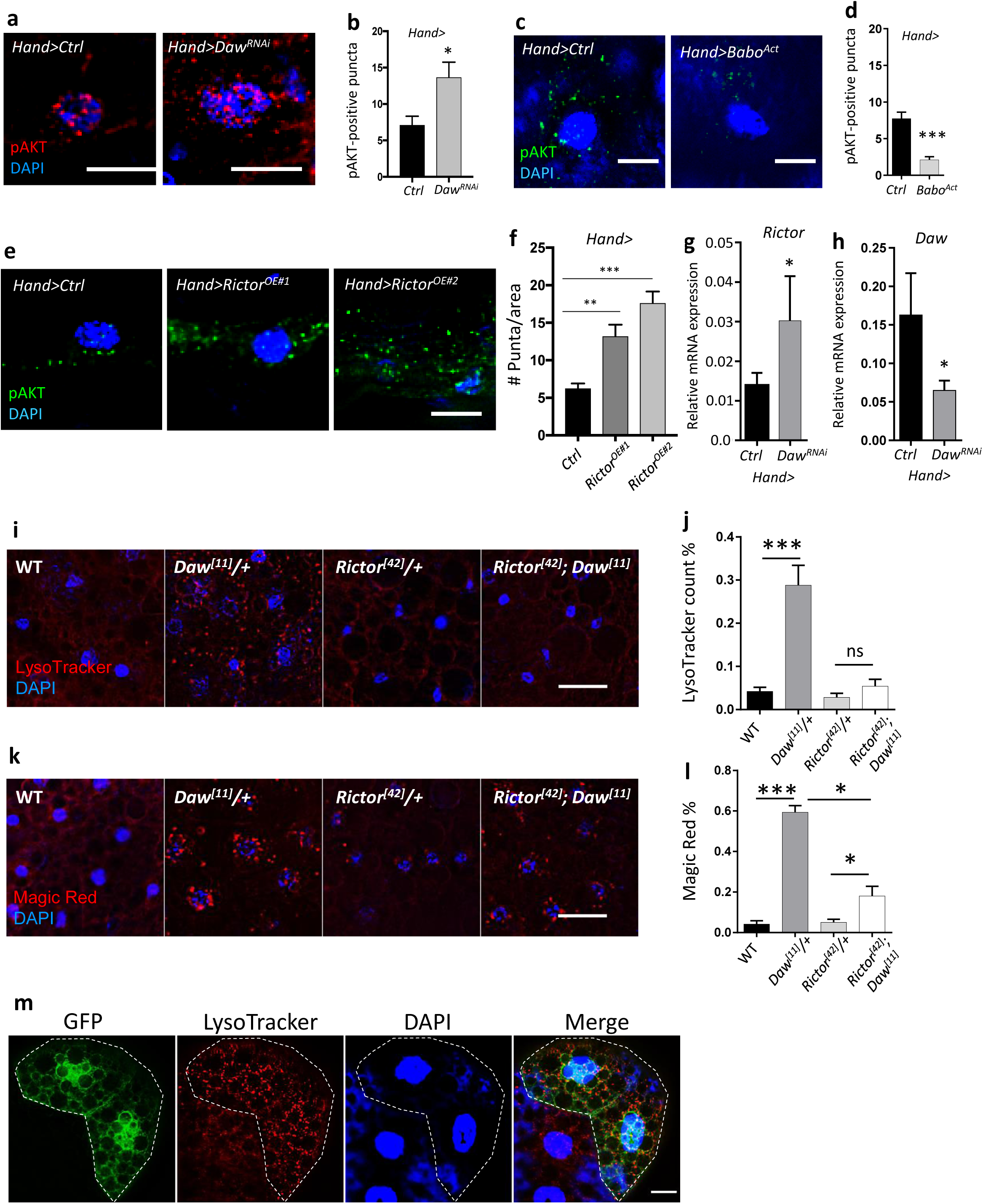
Daw genetically interacts with mTORC2/Rictor to control autophagy. **a.** Representative images of phospho-AKT staining in cardiomyocytes of control and Daw knockdown flies (*Hand-gal4*). Scale bar: 10 μm. **b.** Image quantification of Figure 5a. Student t-test (* p<0.05, ** p<0.01, *** p<0.001). N=5. **c.** Representative images of phospho-AKT staining in cardiomyocytes of control and Daw knockdown flies (*Hand-gal4*). Scale bar: 10 μm. **d.** Image quantification of Figure 5c. Student t-test (* p<0.05, ** p<0.01, *** p<0.001). N=5. **e.** Representative images of phospho-AKT staining in cardiomyocytes of control and Rictor over-expression (*Hand-gal4*). Two independent Rictor over-expression used (#1: *UAS-Rictor*, #2: *UAS-HA-Rictor* ^77^). Scale bar: 10 μm. **f.** Image quantification of Figure 5e. One-way ANOVA (*** p<0.001, ** p<0.01, * p<0.05, ns = not significant). N=5. **g.** QRT-PCR analysis of *Rictor* expression in *Daw* knockdown flies (*Hand-gal4*). Student t-test (* p<0.05, ** p<0.01, *** p<0.001). N=3. **h.** QRT-PCR analysis of *Daw* expression in *Daw* knockdown flies (*Hand-gal4*). Student t-test (* p<0.05, ** p<0.01, *** p<0.001). N=3. **i.** Representative images of LysoTracker staining in adult fat body of *WT*, *Daw*^*[11]*^, *Rictor*^*[42]*^and double mutant *Rictor*^*[42]*^; *Daw*^*[11]*^. Scale bar: 20 μm. **j.** Image quantification of Figure 5i. One-way ANOVA (*** p<0.001, ** p<0.01, * p<0.05, ns = not significant). N=5. **k.** Representative images of Magic Red staining in adult fat body of *WT*, *Daw*^*[11]*^, *Rictor*^*[42]*^*and* double mutant *Rictor*^*[42]*^; *Daw*^*[11]*^. Scale bar: 20 μm. **l.** Image quantification of Figure 5k. One-way ANOVA (*** p<0.001, ** p<0.01, * p<0.05, ns = not significant). N=5. **m.** Representative images of mosaic analysis of LysoTracker staining in larval fat body. Fat body clones were generated by crossing *Rictor* overexpression lines into a FLPout line (*yw*, *hs-flp*, *UAS-CD8::GFP; Act>y+>Gal4,UAS-GFP.nls;UAS-Dcr2*). Clones with *Rictor* overexpression are GFP-positive cells (dashed lines). Scale bar: 20 μm.

Unlike mTORC1, the role of mTORC2 in the regulation of autophagy is unknown. Intriguingly, we found that loss-of-function mutation in *Rictor* repressed the high lysosomal activities in *Daw*^*[11]*^ mutants (Figs. 5i, 5j, 5k, 5l). Similarly, mutations in Sin1, another core component of mTORC2, blocked the elevated lysosomal activity in *Daw* mutants. Furthermore, mosaic analysis revealed that over-expression of Rictor induced lysosome activity (Fig. 5m). Interestingly, unlike *Tsc1*^*[12]*^ mutants, *Rictor*^*[42]*^ mutants did not repress starvation-induced lysosome activity (Fig. S5c). Similarly, activated Babo also did not block starvation-induced lysosome activity (Fig. S5d). These data suggest that activin signaling and mTORC2/Rictor regulate autophagy differently from mTORC1, and Rictor acts downstream of activin signaling in the regulation of autophagy.

Given that Rictor positively regulates autophagy, we predict that Rictor is a cardiac protective factor. Indeed, cardiac-specific over-expression of *Rictor* blocked age-related increases in diastolic intervals and arrhythmia (Figs. 6a, 6b), while knockdown of *Rictor* induced arrhythmia, especially at old ages (Fig. 6c). Interestingly, over-expression of *Rictor* attenuated the elevated diastolic intervals and arrhythmia in fly hearts with constitutively activated *Babo* (Figs. 6d, 6e), suggesting that fly activin signaling regulates cardiac aging through mTORC2/Rictor (Fig. 6f).

**Figure 6.**
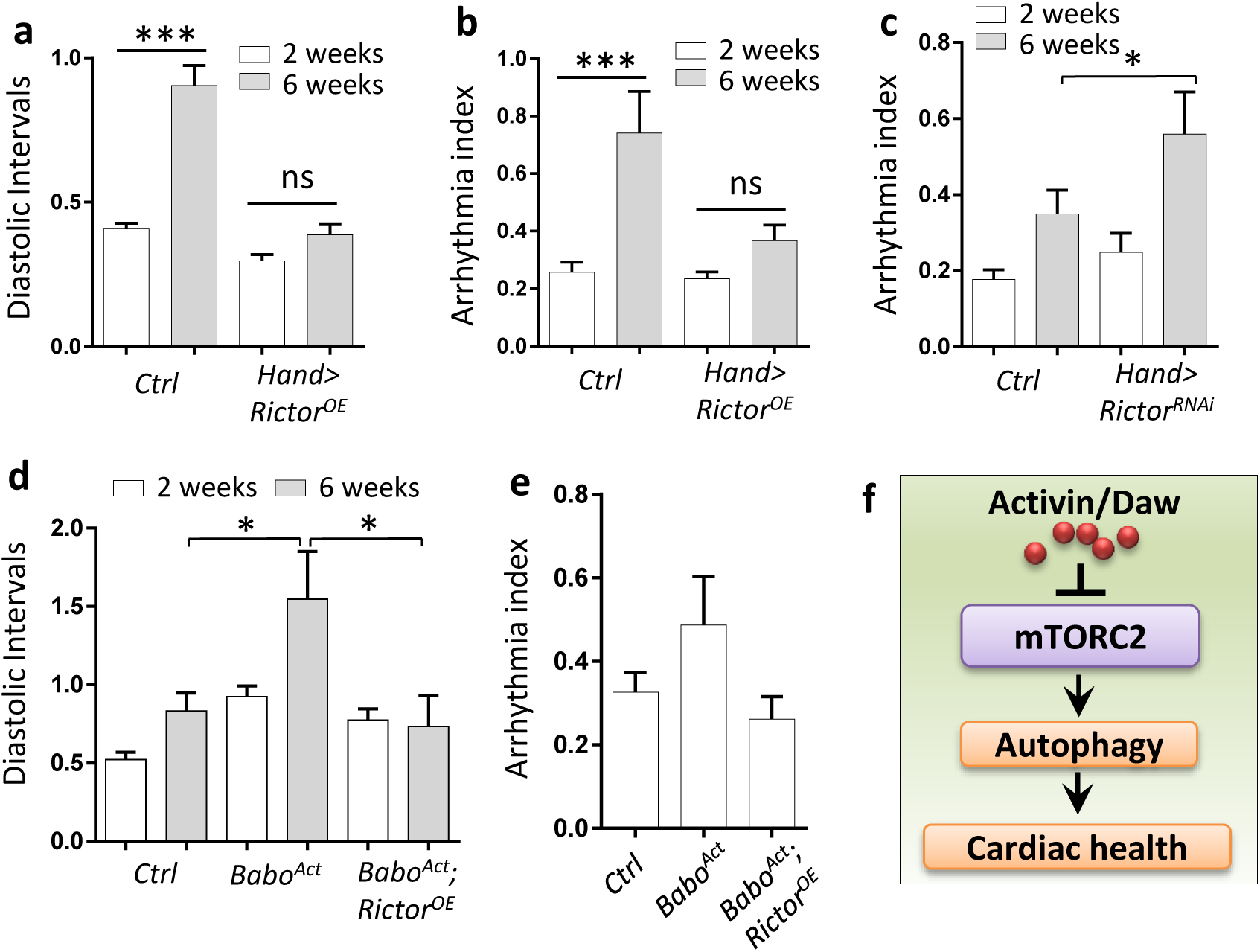
Heart-specific overexpression of *Rictor* preserved cardiac function during aging. **a-b.** Diastolic intervals and arrhythmia in flies with heart-specific overexpression of *Rictor (Hand-gal4*). One-way ANOVA (*** p<0.001, ** p<0.01, * p<0.05, ns = not significant). N=19~26. **c.** Cardiac arrhythmia in flies with heart-specific knockdown of *Rictor*. One-way ANOVA (*** p<0.001, ** p<0.01, * p<0.05, ns = not significant). N=24. **d-e.** Diastolic intervals and arrhythmia in flies with heart-specific expression of *Babo*^*Act*^, or both *Babo*^*Act*^ and *Rictor (Hand-gal4*). One-way ANOVA (*** p<0.001, ** p<0.01, * p<0.05, ns = not significant). N=11~27. **f.** Proposed model for activin-mTORC2 interaction in the regulation of autophagy and cardiac aging.

### Cardiac-specific reduction of activin-like protein *Daw* prolongs lifespan

Tissue-specific hormonal signaling in systemic aging control have been previously reported ^8, 44, 45^. However, whether longevity can be achieved by maintaining healthy hearts remains unclear. Here we tested the effects of cardiac-specific knockdown of activin signaling in longevity control. Knockdown of *Daw* in the heart using two cardiac tissue drivers (*Hand-gal4* and *TinC-gal4*) significantly prolongs lifespan (range from 14% to 32% extension of median lifespan) (Figs. 7a-7e). Compared to *Hand-gal4*, cardiomyocyte-specific driver *TinC-gal4* produced stronger lifespan extension. Two independent lifespan analyses were performed and both showed similar lifespan phenotypes (Fig. 7e). In the first lifespan trial, multiple control lines (dashed lines in Fig. 7a) and two Daw RNAi lines (solid lines in Fig. 7a) were included. Because Daw is a hormonal factor, it is unclear whether Daw regulates lifespan through cell-autonomous or non-autonomous mechanisms. However, our previous study showed that Daw regulates longevity in a tissue-specific manner ^21^. Flight muscle-specific knockdown of *Daw* prolongs lifespan, while fat body-specific knockdown of *Daw* shortens lifespan. Therefore, it is possible that cardiac-specific Daw signaling regulates longevity through maintenance of cardiac health and production of unknown systemic factors from the fly heart.

**Figure 7.**
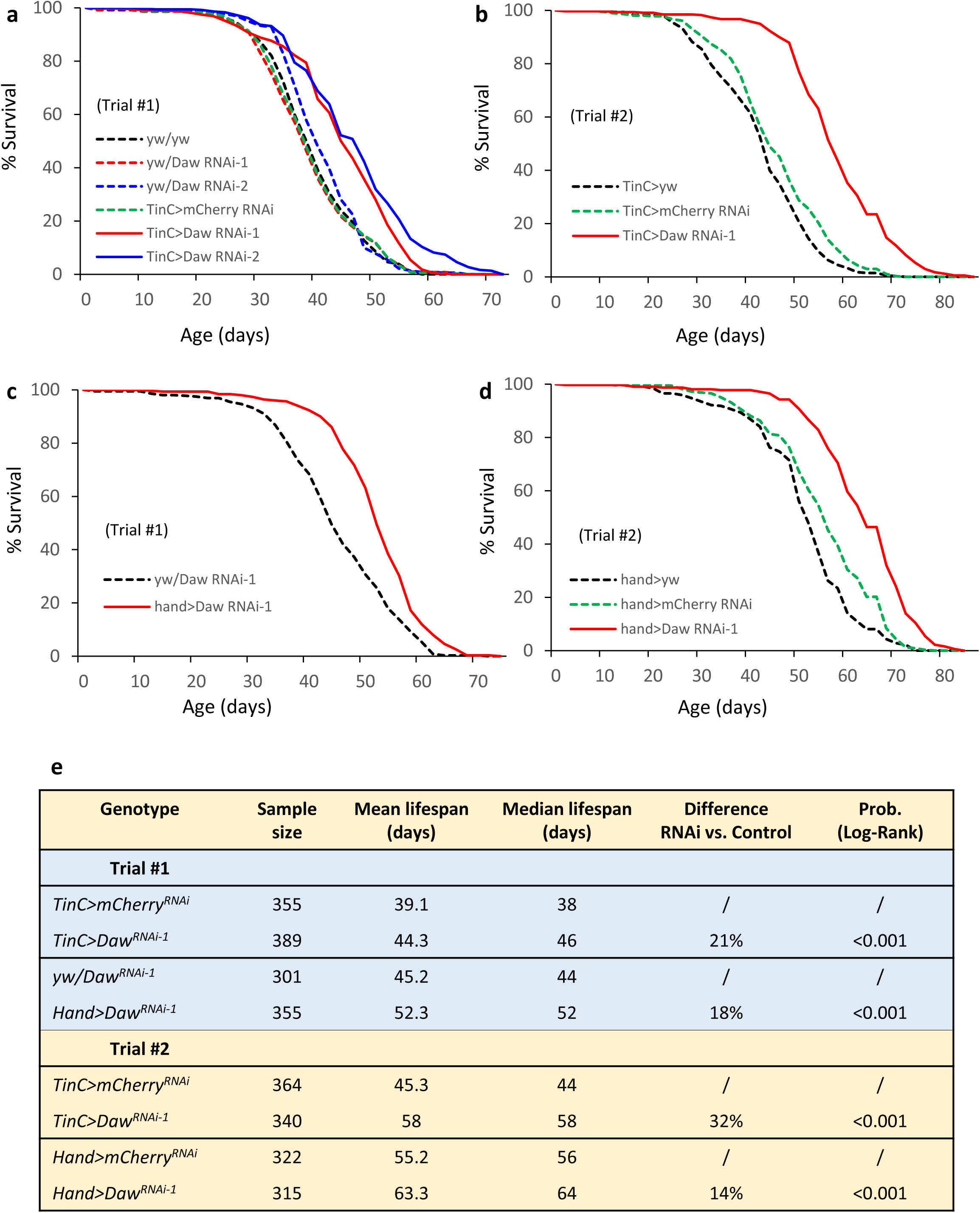
Cardiac-specific knockdown of *Daw* prolongs lifespan. **a-b.** Survival analysis of cardiomyocyte-specific (*TinC-gal4*) knockdown of *Daw*. Two independent trials and multiple controls (*yw x yw*, *yw x Daw RNAi*, *TinC x mCherry RNAi*) were performed. Log-Rank test, p<0.001. **c-d.** Survival analysis of heart-specific (*Hand-gal4*) knockdown of *Daw*. Two independent trials were performed. Log-Rank test, p<0.001. **e.** Lifespan table to show sample size and median lifespan of the survival analysis.

## Discussion

TGF-beta signaling plays vital roles in a wide range of human diseases, including cancer and cardiovascular diseases ^46^. Originally identified as a reproductive hormone, activin, a TGF-beta subfamily member, has become an emerging target for the treatment of many human diseases ^47^. In the present study, we investigated the role of activin in cardiac aging using a *Drosophila* heart model. Similar to our previous study in *Drosophila* flight muscle ^21^, reduction of activin signaling prevented age-related decline of cardiac function. We think that this is partly due to the activation of autophagy. *Daw* mutant flies exhibited high autophagic flux and elevated lysosome activities. Genetic analysis further showed that autophagy gene Atg1 is required for Daw-regulated cardiac aging. Importantly, we found that activation of mTORC1 did not rescue the autophagy and cardiac aging phenotypes in *Daw* knockdown flies. Instead, activin/Daw signaling interacts with mTORC2 subunit Rictor to regulate autophagy and cardiac function. Thus, our studies uncover an emerging role of activin signaling and mTORC2 in the regulation of autophagy and cardiac tissue homeostasis.

Although previous studies reveal an interesting link between TGF-beta family proteins and animal aging ^21, 48-50^, the role of activin in cardiac aging is not well understood. It’s known that activin A serum levels significantly increase with age ^51^. The mRNA expression of *activin A*, as well as serum activin A levels are positively correlated with heart failure ^52, 53^, and hypertension in the elderly ^54^. These findings suggest that activation of activin signaling at old ages might be detrimental to the heart. Studies showed that activin A promotes cardiac fibrosis and myocardial damage after ischemia reperfusion ^55^. Consistently, mice with heterozygous mutations of ALK4 are protected from pressure overload-induced fibrosis and dysfunction ^26, 27^. Inhibition of ALK5 also reduced myocardial infarction-induced systolic dysfunction and left ventricular remodeling in rats ^28^. However, other studies suggest that activin A protects hearts from hypoxia/reoxygenation‐ and ischemia/reperfusion-induced cell death ^29^, even though it can promote cardiac apoptosis at higher concentrations ^56^. The findings from the present study support the negative regulatory role of activin signaling in cardiac function, in particular age-related cardiac dysfunction (e.g., arrhythmia). This is consistent with the pro-aging role of TGF-beta and activin ^21, 50^.

Activin signaling has been implicated in the regulation of a wide range of cellular processes, including cell death and proliferation, inflammation, fibrosis, and metabolic homeostasis ^47, 57-59^. The role of activin signaling in the control of autophagy, a key tissue maintenance process, was only identified recently ^21, 60-63^. In *C. elegans*, reduction of activin-like protein Daf-7 suppressed beta-amyloid peptide-induced autophagosome accumulation ^62, 63^. Another recent study found that activin A blocked oxygen-glucose deprivation-induced autophagy through the inhibition of JNK and p38 MAPK pathways in neuronal PC12 cells ^60^. Our previous study showed that knockdown of *Drosophila* activin-like protein *Daw* in flight muscle induced autophagosome formation and autophagy gene expression ^21^. However, how activin signaling regulates autophagy and how this regulation contributes to tissue homeostasis during aging remain largely unknown. Our findings in the present study suggest that the fly activin-like protein Daw promotes cardiac aging through inhibition of autophagy. It seems that activin signaling controls both autophagosome formation and turnover. Importantly, fly hearts with reduced *Daw* expression maintain high levels of autophagosome turnover (autophagic flux) throughout the life, whereas the autophagosome turnover is largely blocked in aged hearts in wild-type flies. The active autophagosome turnover (and degradation capacity) at old ages is likely responsible for the healthy hearts seen in *Daw* knockdown flies.

mTORC1 is a well-known autophagy regulator and it inhibits autophagosome formation by increasing the phosphorylation of the serine/threonine protein kinase Atg1 (ULK1/ULK2 in mammals), the initiator of the autophagy machinery ^7^. Intriguingly, activation of mTORC1 did not block the induction of autophagosome formation in *Daw* knockdown flies, suggesting that there might be unknown mTORC1-independent mechanisms involved in Daw-regulated autophagy and cardiac aging. mTORC1-independent autophagy regulation has been observed in several recent studies ^64-66^. A genome-wide screen study identified an important role of type III PI3 kinases in autophagy control under normal nutritional conditions ^64^. The screen also revealed that growth factor signaling pathways, such as MAPK-ERK and AKT/FOXO3, negatively regulate autophagy by inhibiting the type III PI3 kinase cascade. Interestingly, we noticed that knockdown of *Daw* led to elevated AKT phosphorylation and the expression of mTORC2 subunit *Rictor*, suggesting mTORC2/AKT signaling may be involved in Daw-regulated autophagy and cardiac aging. Indeed, we found that mutation of *Rictor* blocked the activation of autophagy in *Daw* knockdown flies. mTORC2 is one of the two mTOR complexes that is insensitive to short-term rapamycin treatment, and is involved in glucose homeostasis and actin cytoskeleton reorganization ^67, 68^. The direct regulatory role of mTORC2/Rictor in cellular autophagy has not been previously established, although a few studies suggest that Rictor may be involved in autophagy activation under unique circumstances. For example, Rictor is required for resveratrol-induced autophagy in rat myocardium and H9c2 cardiac myoblast cells ^69^. In rat kidney NRK-52E cells, silencing Rictor blocked cisplatin induced autophagy ^70^. Our findings align with these studies suggesting that Rictor is a positive regulator of autophagy. However, conflicting findings are reported in other studies showing Rictor negatively regulates autophagosome formation in mouse skeletal muscle ^71^, and LC3-II levels in human senescent endothelial cells ^72^. Thus, more studies are needed to better understand the role of mTORC2/Rictor in the regulation of autophagy and its contribution to tissue homeostasis during aging.

The positive role of Rictor in autophagy control identified in the present study is consistent with the cardioprotection effects of Rictor. We found that cardiac-specific overexpression of *Rictor* slows cardiac aging, similar to *Daw* knockdown. Our findings are consistent with two recent studies showing that disruption of mammalian Rictor induced cardiac dysfunction ^73, 74^. Cardiac-specific deletion of Rictor in mice induced dilation and decreased fractional shortening by activating MST1 kinase and hippo pathway ^74^. Unlike mTORCl mutants, Rictor null mutants (in both worms and mice) are short-lived ^75, 76^, suggesting the detrimental role of mTORC2/Rictor in regulating tissue aging. The role of *Drosophila* Rictor in longevity regulation is not yet known, although it has been shown that *Drosophila* mutants of Rictor and Sin1 decreased the tolerance to heat stress ^77^ and over-expression of *Rictor* rescued Pink1 knockdown-induced mitochondrial aggregation in flight muscle ^78^. The Rictor-Pink1 interaction suggests that Rictor could be a potential positive regulator for mitophagy, a key process for the mitochondria quality control during aging.

During normal aging, the heart undergoes complex phenotypic changes such as progressive myocardial remodeling, reduced myocardial contractile capacity, increased left ventricular wall thickness and chamber size, prolonged diastole as well as increased arrhythmia^3^. All of these biological changes can gradually alter cardiac functions and confer vulnerability of the heart to various cardiovascular stresses, thus increasing the chance of developing cardiovascular disease dramatically. The present study has uncovered an important role of TGF-beta/activin signaling pathway in the regulation of autophagy and cardiac aging in *Drosophila*. We also found an intriguing interaction between activin and mTORC2 in the regulation of cardiac autophagy, as well as a new role of Rictor in cardiac aging in *Drosophila*. It remains to be determined how activin regulates Rictor in fly hearts. It is possible that activin signaling regulates Rictor through both transcriptional activation (e.g., through insulin/FOXO signaling ^79^) and post-translational modification.

Additionally, cardiac-specific knockdown of activin-like protein *Daw* extended fly lifespan, suggesting that longevity can be achieved by maintaining a healthy heart. Our findings also suggest that Daw itself or other unidentified hormonal factors secreted from cardiomyocytes play an important role in systemic aging control. Hence, it remains to be determined the mechanistic basis for activin-mediated control of cardiac aging and longevity, which will eventually help the development of therapeutic interventions targeting activin for the treatment of age-related cardiovascular diseases.

## Materials and methods

### Fly husbandry and stocks

Flies were maintained at 25°C, 60% relative humidity and 12-hour light/dark (40% relative humidity was used in the experiments from Figures 1&2, which were performed in Bodmer laboratory). Adults were reared on agar-based diet with 0.8% cornmeal, 10% sugar, and 2.5% yeast (unless otherwise noted). Fly stocks used in the present study are: *UAS-Daw RNAi* (Bloomington *Drosophila* Stock Center (BDSC) #34974, BDSC #50911, and Vienna Drosophila Resource Center (VDRC) #105309), *UAS-Babo RNAi* (BDSC #25933, BDSC #40866), *UAS-Babo-Act* (also known as *Babo.Q302D*) ^80^, *UAS-Tsc1 RNAi* (BDSC #52931, #54034), *UAS-Reptor RNAi* (BDSC #25983), *UAS-Rheb* (BDSC #9688), *UAS-Atg1 RNAi* (BDSC #26731), *UAS-Rictor RNAi* (BDSC #36699), *Daw*^*[11]* 81^, *Tsc1*^*[12]*, 82^, *Rictor*^*[42]* 83^, *Sin1*^*[e03756]*^ (BDSC #18188), *UAS-Rictor*^77^, *UAS-HA-Rictor*^*77*^, *Hand-gal4* (also known as Hand4.2-gal4) ^32^, and *TinC-gal4* (also known as *TinC.4-gal4*) ^33^, *Dot-gal4* ^34^, *Hand-GS-gal4* ^84^, *UAS-mCherry-Atg8a* (BDSC #37750), *Atg8a*^Δ*4*, 85^.

*UAS-Daw RNAi* (BDSC #34974), *UAS-Babo RNAi* (BDSC #25933), and *UAS-Myo RNAi* (BDSC #31114) were backcrossed into *yw*^*R*^ background for 5~7 generations, and *yw*^*R*^ flies were used as control or wild-type (WT) flies in most of the experiments. For other UAS-RNAi lines that were not backcrossed to *yw*^*R*^, following genotypes were used as control: *y*^*1*^ *sc*^*^ *v*^*1*^; *P[VALIUM20-mCherry]attP2* (BDSC #35785), *y*^*1*^ *v*^*1*^; *P[CaryP]attP2* (BDSC #36303), or *y*^*1*^ *v*^*1*^; *P[CaryP]attP40* (BDSC # 36304). Female flies were used in all experiments. RU486 (mifepristone, Sigma, St. Louis, MO, USA) were used to activate *Hand-GS-Gal4* at a final concentration of 200 μM mixed in food.

### Mosaic analysis

Two FLPout lines were used: 1). *hs-flp; endogenous P-3x mCherry-Atg8a*, *UAS-GFP/Cyo; Act>CD2>Gal4,UAS-Dcr2/TM6*) (kindly provided by Gábor Juhász). 2).*yw, hs-flp, UAS-CD8::GFP; Act>y+>Gal4,UAS-GFP.nls;UAS-Dcr2/SM6::TM6B* (kindly provided by Jun-yuan Ji, originally generated by Bruce Edgar). To generate clones, freshly laid eggs (within 4-6 hr) were heated in a 37 °C water bath for 45 min. Early L3 larvae (84 hours after egg laying) were used in the experiments.

### Fly heartbeat analysis

To measure cardiac function parameters, semi-intact *Drosophila* adult fly hearts were prepared according to previously described protocols ^31^. High-speed 3000 frames movies were taken at a rate of 100 frames per second using a Hamamatsu ORCA-Flash4.0 digital CMOS camera (Hamamatsu Photonics, Hamamatsu, Japan) on an Olympus BX51WI microscope with a 10X water immersion lens (Olympus, Waltham, MA, USA) (Hamamatsu EM-CCD 9300 camera was used in the experiments from Figures 1&2, which were performed in Bodmer laboratory). The live images that contain heart tub within abdominal A3 segment were processed using HCI imaging software (Hamamatsu Photonics, Hamamatsu, Japan). M-modes and cardiac parameters were generated using SOHA, a MATLAB-based image application as described previously ^31, 86^. The M-mode provides a snapshot of movement of heart wall over time. Four cardiac parameters were analyzed, including heart period, diastolic interval, systolic interval, and arrhythmia index. Diastolic interval (DI) is the duration for each heart relaxation phase (diastole). Heart period (HP) is the pause time between the two consecutive diastole (HP is the reciprocal of heart rate). Systolic interval (SI) was calculated as the HP minus the DI. Arrhythmia index (AI) is the standard deviation of all HP in each fly normalized to the median HP. Cardiac output was calculated using following equation: (π r(d)^2^ - π r(s)^2^) x HR. r(d) is radius of the heart tube at diastolic phase, while r(d) is radius of the heart tube at systolic phase.

### Quantitative RT-PCR

RNA extraction and cDNA synthesis were performed using Cells-to-CT kit (Thermo Fisher Scientific, Waltham, MA, USA) from ~15 dissected adult hearts. QRT-PCR was performed with a Quantstudio 3 Real-Time PCR System (Thermo Fisher Scientific, Waltham, MA, USA). mRNA abundance of each gene was normalized to the expression of ribosomal protein L32 (*RpL32* or *rp49*) by the method of comparative C_T_. Primer sequences are listed in Supplementary Table S1.

### Antibodies and immunostaining

Since commercial antibodies for *Drosophila* Atg8a show non-specific staining in immunostaining of fly tissues (data not shown), we used a GABARAP (E1J4E) antibody (1:300) (Cell Signaling Technology (CST) #13733, Danvers, MA, USA) to stain and detect endogenous *Drosophila* Atg8a. The epitope used to generate GABARAP antibody has identical amino acid sequences to *Drosophila* Atg8a protein (personal communication with Cell Signaling Technology, and see sequence alignment in Fig. S1a). This antibody has been previously used in several *Drosophila* autophagy studies ^87-89^, and verified by Kim et al. ^90^. We further verified the specificity of the GABARAP antibody using an Atg8a loss-of-function mutant *Atg8a*^Δ*4*^ (Figs. S1b-S1d). Other antibodies used for immunostaining are list below: Ref(2)P (1:1000) ^38^, Phospho-Smad2 antibody (1:500) (CST #3108), Phospho-Akt (Ser473) (1:300) (CST # 4060), Alexa Fluor 488-conjugated Phalloidin (or Alexa Fluor 594) for F-actin staining (Thermo Fisher Scientific, Waltham, MA, USA). All fluorescence-conjugated secondary antibodies were from Jackson ImmunoResearch, West Grove, PA, USA).

For immunostaining, adult female flies were collected and dissected in PBS. Hearts were fixed in 4% paraformaldehyde for 15 min at room temperature (RT). After washing in PBS with 0. 1% Triton X-100 (PBST), the fixed hearted were blocked in 5% normal goat serum (NGS) diluted in PBST for 1 hour at RT. Hearts were then washed with PBST and incubated overnight at 4°C with primary antibodies diluted in 5% NGS. After washing with PBST, the samples were incubated for 2 hours at RT with appropriate fluorescence-conjugated secondary antibodies.

Hearts were mounted in ProLong Gold anti-fade reagent (Thermo Fisher Scientific, Waltham, MA, USA) before imaged using an epifluorescence-equipped BX51WI microscope (Olympus, Waltham, MA, USA).

### Imaging analysis

For imaging analysis, fluorescence images were first processed using the deconvolution module in Olympus cellSens Demensions software, and then the number of puncta in a selected perinuclear region (~707 μm^2^) was measured with the “Measure and Count” module in cellSens software (Olympus, Waltham, MA, USA).

### LysoTracker and Magic Red staining

Acidic organelles (including autolysosome) were monitored by staining tissue with 100 nM LysoTracker Red DND-99 (Invitrogen, Grand Island, NY, USA) for 5 minutes at room temperature. Lysosomal cathepsin B activities were monitored using Magic Red Cathepsin-B Assay kit (#938, ImmunoChemistry Technologies, Bloomington, MN, USA) following manufacturer’s manual. Nuclei were stained with either DAPI or Hoechst 33342 (1 μg/ml) (ImmunoChemistry Technologies).

### Western blotting for Atg8a lipidation

Antibodies for western blot included: β-actin antibody (1:2000) (CST #4967), GABARAP (E1J4E) antibody (1:2000) (CST #13733), and HRP-conjugated secondary antibodies (Jackson ImmunoResearch, West Grove, PA, USA). Flies were first transferred to centrifuge tube containing 1x laemmli buffer (10 μl buffer for each milligram of fly), and heated at 100°C for 3 min. The samples were then homogenized and heated again at 100°C for 3 min. After the centrifuge at 13000x rpm for 5 min, supernatants were collected and loaded onto Mini-PROTEAN precast gels (Bio-Rad, Hercules, CA, USA). Following incubation with primary and secondary antibodies, the blots were visualized with Pierce ECL Western Blotting Substrate (Thermo Fisher Scientific, Waltham, MA, USA).

### Bafilomycin A1 and Chloroquine Treatment

For Bafilomycin A1 treatment, semi-intact hearts were incubated with 100 nM of bafilomycin A1 (Alfa Aesar, Haverhill, MA, USA) in artificial hemolymph (buffer receipt in ^31^) for two hours at room temperature prior to the appropriate immunostaining. DMSO was used as a control.

For chloroquine treatments, 100 μl of 20 mM chloroquine diphosphate salt (Sigma-Aldrich, St Louis, MO, USA) was added onto the fly food. Flies were fed with chloroquine for at least 24 hours prior to the cardiac analysis or the western blots.

### Demography and survival analysis

Newly enclosed female flies were allowed to mate for two days, then separated from males and assigned to replicate one-liter demography cages at a density of 100-125 flies per cage. Three independent cages were set-up per genotype. Food was changed every two days, at which time dead flies were removed from the cage and counted. Survival analysis was conducted with JMP statistical software (SAS Institute, Cary, NC, USA), and data from replicate cages were combined. Mortality distributions were compared by Log-rank test.

### Statistical analysis

GraphPad Prism (GraphPad Software, La Jolla, CA) was used for statistical analysis. To compare the mean value of treatment groups versus that of control, either student t-test or oneway ANOVA was performed using Dunnett’s test for multiple comparison. The effects of mutants during aging was analyzed by two-way ANOVA, including Tukey multiple comparisons test. In SOHA analysis, the outliers were identified using Robust regression and Outlier removal (ROUT) method (Q=1%) prior to the data analysis.

## Acknowledgements

We thank Bloomington Drosophila Stock Center, Harvard Medical School, and Drosophila Genomics Resource Center for fly stocks and cDNA clones. We thank Michael O’Connor for activin reagents and fly stocks, Duojia Pan, Stocker Hugo, Jongkyeong Chung, Gábor Juhász, Jun-yuan Ji, Bruce Edgar for fly stocks. Marc Tatar for help with Demography and survival analysis. This work was supported by NIH/NIA R00 AG048016 and AFAR Research Grants to HB.

## Author Contributions Statement

Conceived and designed the experiments: KC RB KO HB. Performed the experiments: KC PK YL KH APS IPN HB. Analyzed the data: KC PK YL KH ET APS IPN HB. Wrote the paper: KC RB KO HB. All authors reviewed and approved the manuscript.

## Supplemental file

**Table S1.**
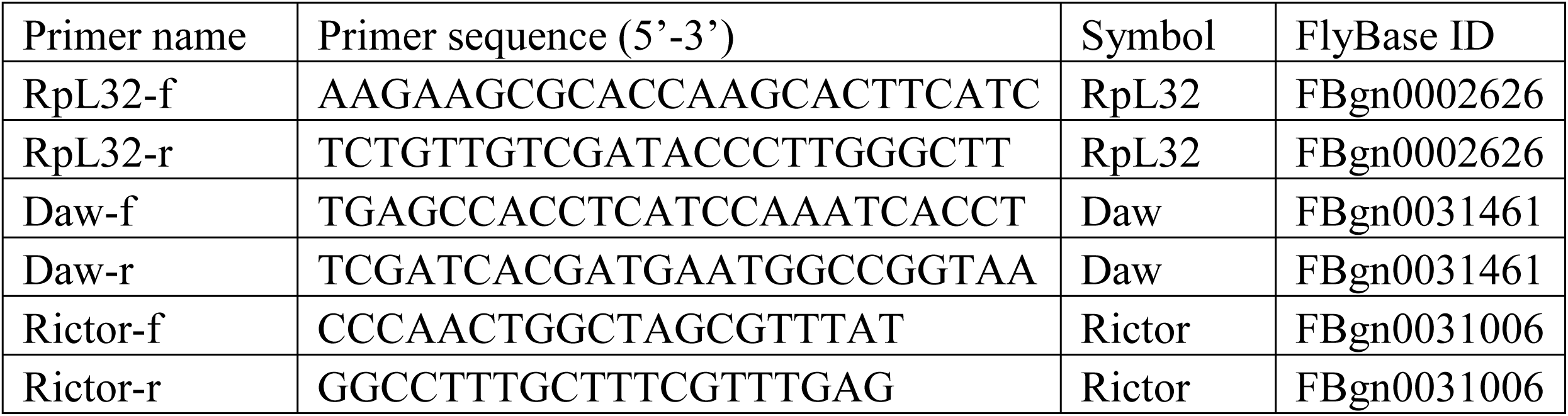
Primer list

**Figure S1.**
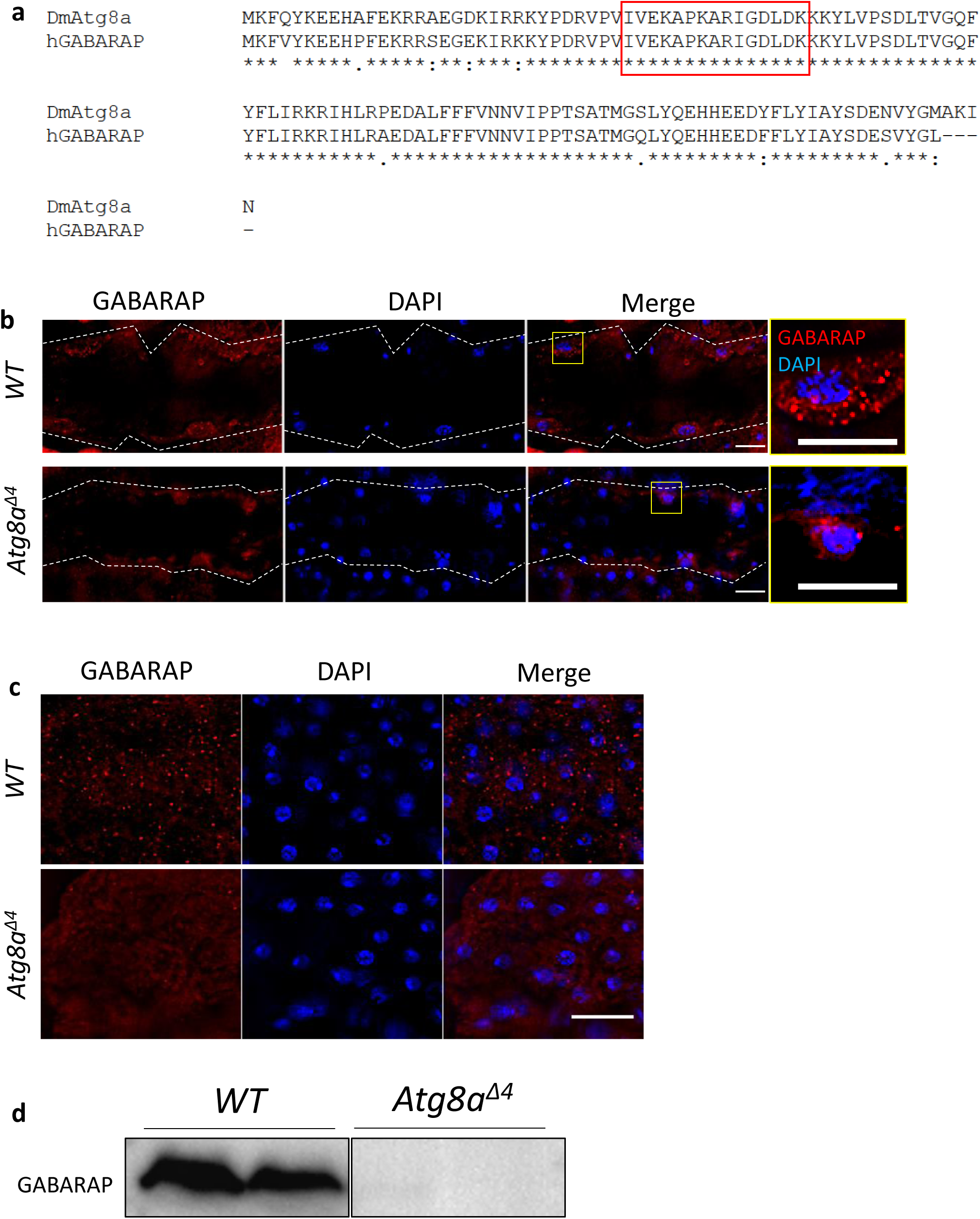
Validation of GABARAP antibody. **a.** Multiple sequence alignment between *Drosophila* Atg8a and human GABARAP using ClustalW (https://www.genome.jp/tools-bin/clustalw). The red box indicates the predicted epitope sequences used in producing GABARAP antibody. **b.** Representative images of GABARAP immunostaining in fly hearts of *WT* and *Atg8a*^Δ*4*^ mutants. Heart tube is located between two yellow dashed lines. The panels on the right are the zoomed-in images. Scale bar: 20 μm. **c.** Representative images of GABARAP immunostaining in adult fat body of *WT* and *Atg8a*^Δ*4*^ mutants. The panels on the right are the zoomed-in images. Scale bar: 20 μm. **d.** Western blots testing GABARAP antibody using *Atg8a*^Δ*4*^ mutants.

**Figure S2.**
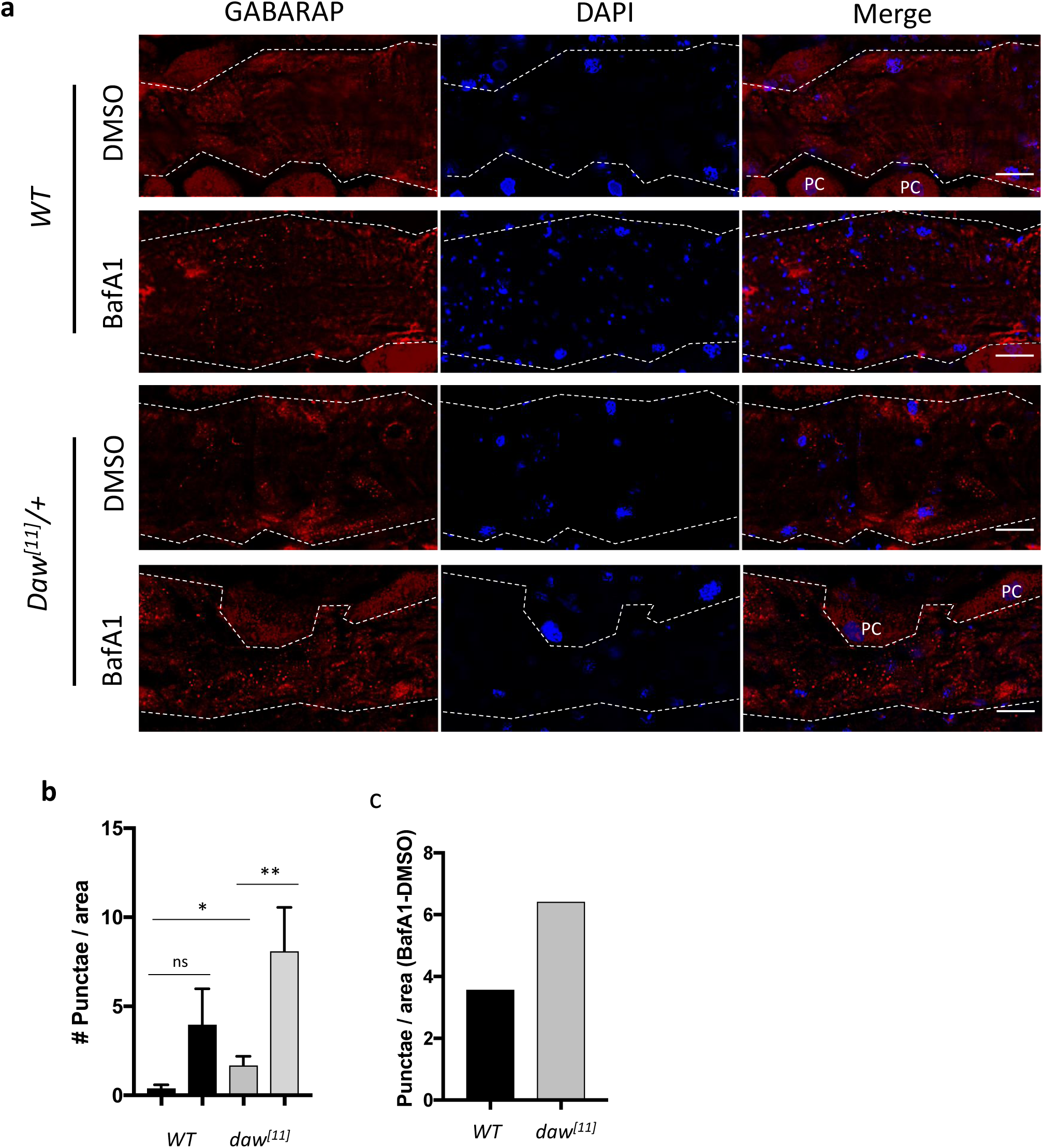
Daw inhibits autophagy flux in the heart. **a.** Representative images of GABARAP immunostaining in fly hearts of *WT* and *Daw*^*[11]*^ mutants treated with or without Baf A1. Heart tube is located between two yellow dashed lines. The panels on the right are the zoomed-in images. PC: Pericardial cell. Scale bar: 20 μm. **b.** Quantification of Figure S2. One-way ANOVA (*** p<0.001, ** p<0.01, * p<0.05, ns = not significant). N=5. **c.** Quantification of Baf A1-induced puncta between *WT* and *Daw*^*[11]*^ mutants.

**Figure S3.**
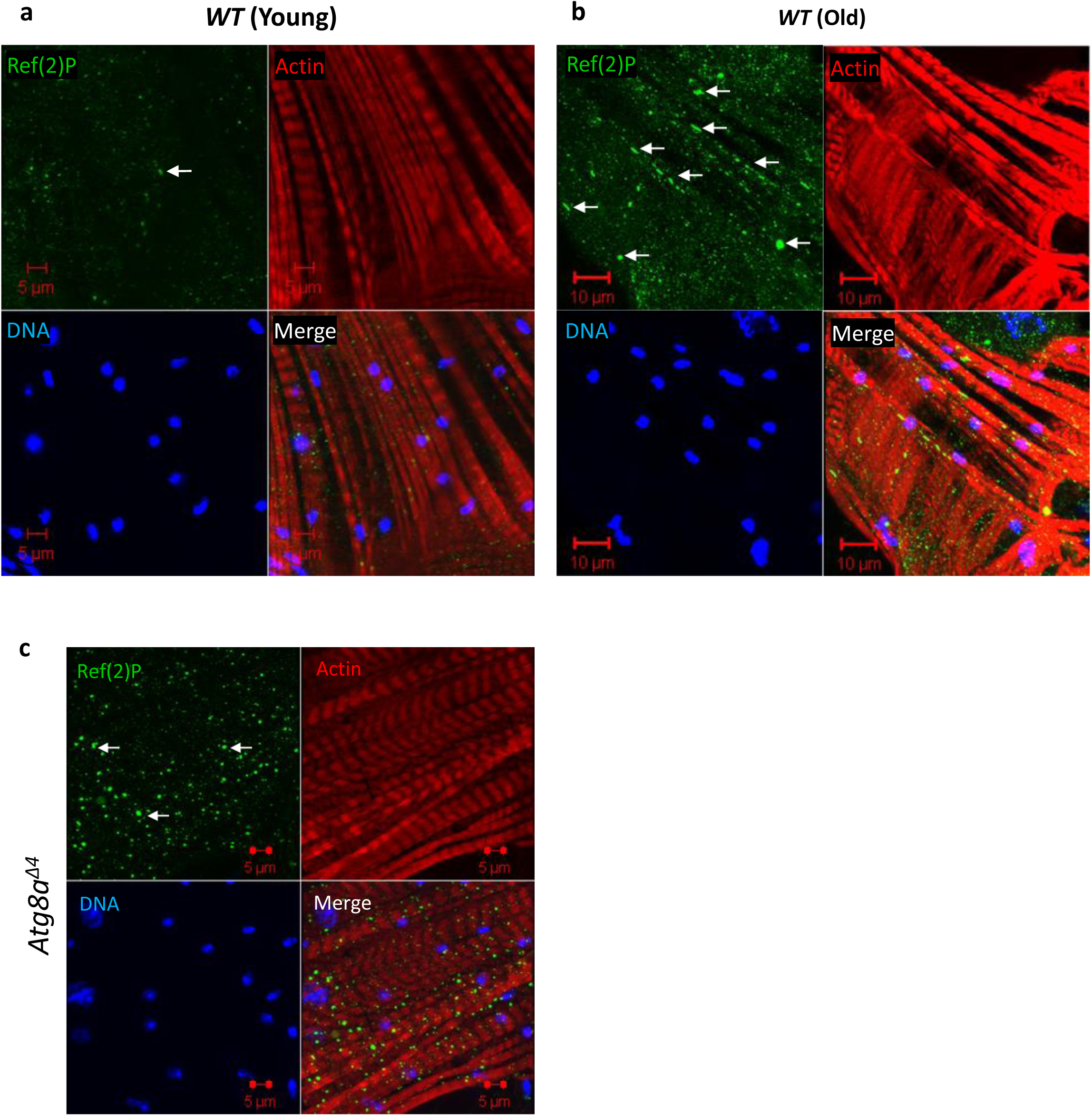
Aging increases the accumulation of Ref(2)P in the heart. **a-b.** Representative images of Ref(2)P in young and old hearts. Scale bar: 10 μm. **c.** Representative images of Ref(2)P in the heart of *Atg8a*^Δ*4*^ mutants. Scale bar: 10 μm.

**Figure S4.**
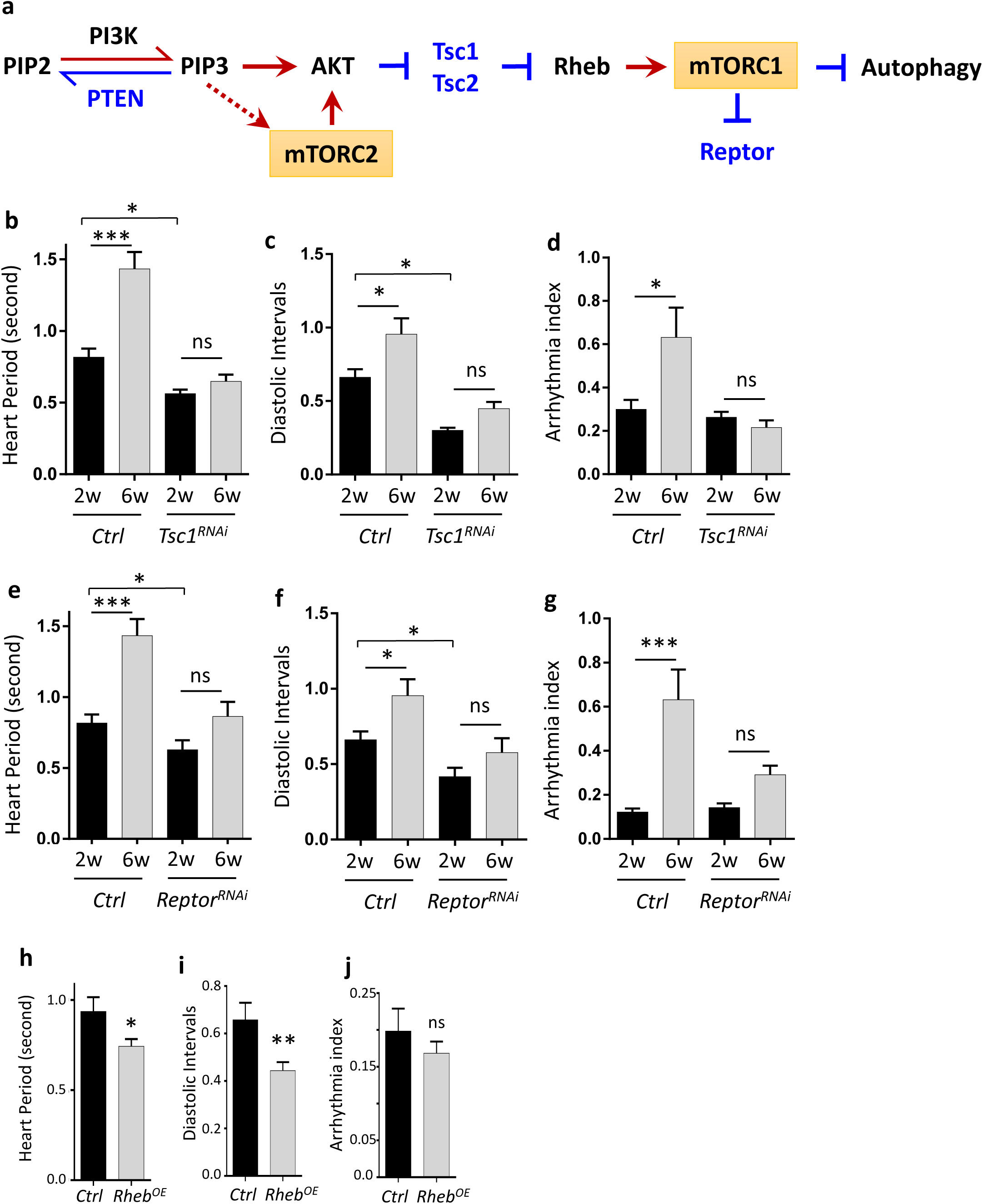
Activation of mTORC1 alters heart period, but not arrhythmia. **a.** Schematic diagram for PI3K/AKT/mTOR signaling pathway. **b-d.** Heart period, diastolic intervals, and arrhythmia in control (*Ctrl*) and cardiomyocyte-specific *Tsc1* knockdown flies (*TinC-gal4*). One-way ANOVA (*** p<0.001, ** p<0.01, * p<0.05, ns = not significant). N=14~18. **e-g.** Heart period, diastolic intervals, and arrhythmia in control (*Ctrl*) and cardiomyocyte-specific *Reptor* knockdown flies (*TinC-gal4*). One-way ANOVA (*** p<0.001, ** p<0.01, * p<0.05, ns = not significant). N=13~28. **h-j.** Heart period, diastolic intervals, and arrhythmia in control (*Ctrl*) and flies overexpressing *Rheb (TinC-gal4*). One-way ANOVA (*** p<0.001, ** p<0.01, * p<0.05, ns = not significant). N=21~31.

**Figure S5.**
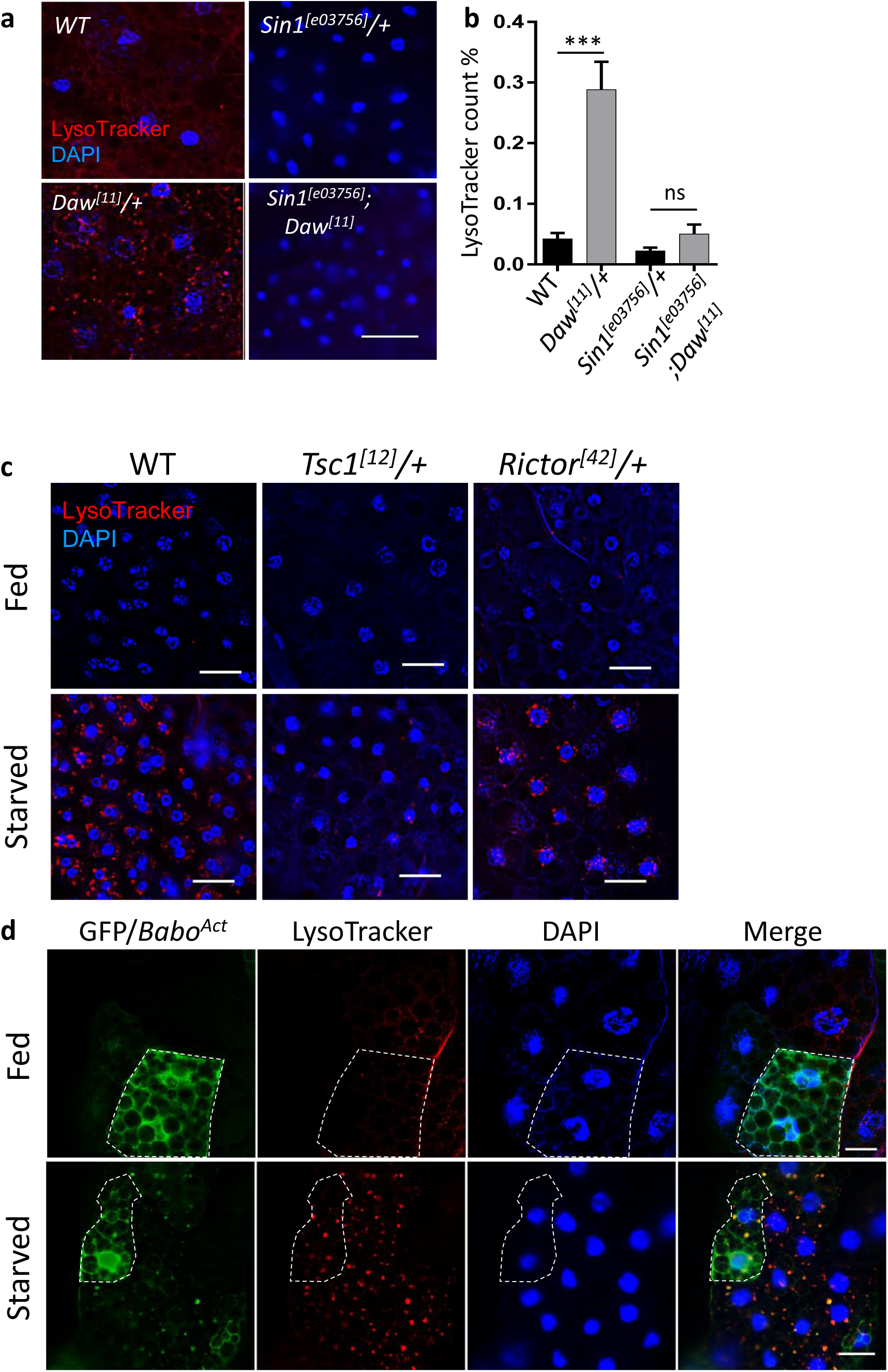
Autophagy regulation by mTORC2. **a.** Representative images of LysoTracker staining in adult fat body of *WT*, *Daw*^*[11]*^, *Sin1*^*[e03756]*^ and double mutant *Sin1*^*[e03756]*^; *Daw*^*[11]*^. Scale bar: 20 μm. **b.** Image quantification of Figure S5a. One-way ANOVA (*** p<0.001, ** p<0.01, * p<0.05, ns = not significant). N=5. **c.** Representative images of LysoTracker staining in fed and starved WT, *Tsc1* mutants, and *Rictor* mutants. Scale bar: 20 μm. **d.** Representative images of mosaic analysis on LysoTracker staining in larval fat body of *Babo*^*Act*^ flies upon starvation. Larval fat body clones were generated using a FLPout line (*yw*, *hsflp*, *UAS-CD8::GFP; Act>y+>Gal4,UAS-GFP.nls;UAS-Dcr2*). Clones with *Babo*^*Ac*^ expression are GFP-positive cells (dashed lines). Scale bar: 20 μm.

